# A high-throughput *in vitro* translation screen towards discovery of novel antimalarial protein translation inhibitors

**DOI:** 10.1101/248740

**Authors:** Hella Baumann, Holly Matthews, Mengqiao Li, Jenny J. Hu, Keith R. Willison, Jake Baum

## Abstract

Drugs that target protein synthesis are well-validated for use as antimicrobials, yet specific high throughput (HTP) methods to screen for those targeting malaria are lacking. Here, we have developed a cell free *in vitro* translation (IVT) assay for the human malaria parasite, *Plasmodium falciparum*, which reconstitutes the native parasite protein translation machinery. Combining clarified IVT lysate with a click beetle luciferase reporter gene fused to untranslated regions of *Pf histidine-rich proteins* (hrp)-2 and 3, the HTP IVT assay accurately reports protein translation in a 384-well plate format using a standard spectrofluorometer. We validate the assay as effective in detecting compounds targeting the ribosome, ribosome co-factors (elongation factor 2) and cytosolic tRNA synthetases as well as its ability to find translation inhibitors in a blind screen using a high-density assay format amenable for high throughput. This demonstrates an ability to reconstitute the breadth of the parasite eukaryotic protein translation machinery *in vitro* and use it as a powerful platform for antimalarial drug discovery.

## INTRODUCTION

Despite marked progress in reducing deaths from malaria, the disease remains a major cause of infant mortality in the developing world, with more than 500,000 estimated to die each year ^1, 2^. In the absence of a fully protective vaccine ^3^, chemotherapeutics constitute the best available clinical tool to prevent or treat malaria disease. The most effective formulations, and standard of care recommended by the World Health Organization (WHO), are based on combination therapies with endoperoxides such as artesunate, artemisinin and artemether, referred to as artemisinin-based combination therapies (ACTs). However, like every clinical antimalarial to date, evidence is growing for the emergence of resistance to ACTs, specifically in South East Asia though not as yet in Africa ^4, 5^. To combat ACT resistance and prevent its potentially devastating spread to sub Saharan Africa new drugs are required with novel modes of action ^6^.

In recent years, cell-based screens have demonstrated great power in their capacity to discover compounds for development as next generation antimalarials. Notably, several of these screens have repeatedly found compounds with activity against malaria parasite protein synthesis ^7-9^. Protein synthesis in the malaria parasite is an attractive target given its essential nature to parasite cell growth and development and the strong precedence for mRNA to protein translation as an antimicrobial target for bacterial infections ^10^. Screening platforms to specifically identify compounds with similar activity against malaria would be a welcome addition to the antimalarial field to increase and differentiate the drug discovery pipeline. Towards this, two studies have devised formulation of a cell free *in vitro* translation (IVT) lysate ^11^ and its development towards a luciferase reporter assay ^12^. This provides a foundation from which inhibitors specific to protein synthesis in the most virulent human malaria parasite, *Plasmodium falciparum*. Whilst some compounds have been screened with this in mind, for example assaying of the Medicines for Malaria Venture Malaria Box ^12^ (400 bioactive compounds with minimal human cytotoxicity profiles), the potential to translate such a platform to blind, high throughput (HTP) screening for compounds targeting the breadth of the protein translation machinery has not been fully realized. To address this, we sought to develop an optimized 384-well-plate assay for HTP screening, applicable to a conventional laboratory spectrofluorometer and validated for its robustness, replicability and capacity to find inhibitors of protein synthesis with diverse modes of action as demonstrated with known translation inhibitors.

## RESULTS AND DISCUSSION

Building on recent efforts to devise a protein translation assay applicable to HTP screening, we first optimized a protocol for large scale, yet reduced budget, protein lysate production, compatible with protein translation. For the production of 3-8 ml lysate, up to 6 liters of a high parasitemia *P. falciparum* 3D7 culture were saponin-treated, with cytosolic extract released via nitrogen cavitation (see MATERIALS AND METHODS for details) (Figure 1A). This yielded on average a lysate with 1000-2000 μg/mL of total RNA as a measure for presence of ribosomal RNA and hence potential lysate activity. We next sought to develop a reporter system compatible with both a time course and endpoint plate-based assay with optimal signal to noise ratio but using reduced amounts of lysate. A parent vector, pHLH-1 ^13^, was used as the foundation for downstream expression plasmids, in which expression of *Photinus pyralis* firefly luciferase (*luc*) occurs under the control of flanking *P. falciparum* regulatory elements from the histidine rich proteins, namely the 5’ promoter element from *hrp3* and 3′ untranslated region (UTR) from *hrp2*. To generate mRNA from this vector *in vitro*, a T7-terminator was introduced after this expression cassette using site-directed mutagenesis, generating the vector pHLH-T7term (referred to as pHLH here for simplicity).

**Figure 1.**
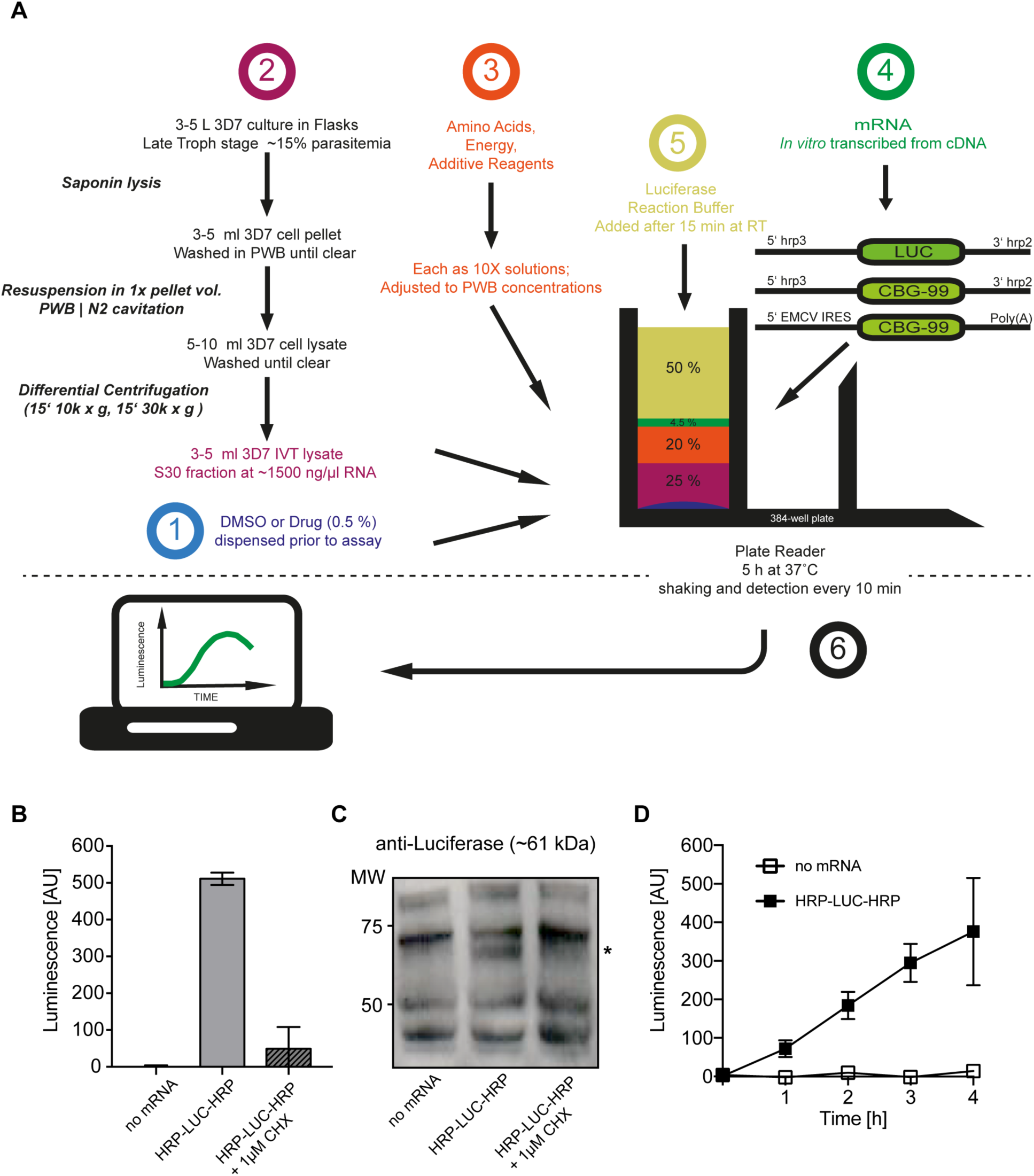
Parasite *in vitro* translation of firefly luciferase (*luc*). **A)** Schematic of IVT lysate production and assay. **B)** Bar diagram demonstrating the *in vitro* translation of *luc* as detected from luminescence after 15 hours incubation. Results are shown as averages of duplicate assays with either DMSO or 1 μM Cycloheximide in DMSO. Error bars show standard deviation. **C)** Demonstration of *luc* expression after 15 hours with western blot. Lysates merged after duplicate assay with either DMSO or 1 μM Cycloheximide in DMSO. Asterisk indicates supposed run length of firefly luciferase **D)** *In vitro* Translation of *luc* over time as measured by luminescence. The graph shows detected averages of triplicate assays with DMSO at indicated time points when luciferase reaction mix was added. Error bars indicate standard deviation.

The mRNA from cDNA was generated by a high yield *in vitro* transcription protocol previously established by Mureev et al ^14^ prior to each assay. Supplementation of the lysate with energy and amino acids demonstrated efficient production of luciferase via pHLH, detectable via luminescence from Oxyluciferin (Figure 1B) or by immunoblot (Figure 1C, asterisk). Adding the translation inhibitor Cycloheximide (CHX) reduced the amount of luciferase detected through luminescence of Oxyluciferin (Figure 1B) and in a western blot (Figure 1C), confirming the protein derives from *in vitro* translation. To validate the functionality of *in vitro* translation reagents and our workflow in a parallel Human embryonic kidney cells 293 (HEK) cell lysate, a further expression vector for *luc* (pIRES-Luc) was generated by cloning the *luc* expression cassette into a pT7CFE vector (see MATERIALS AND METHODS) generating an expression construct in which luciferase is flanked by a 5’ internal ribosomal entry site (IRES) from encephalomyocarditis virus (EMCV) and a 3’ polyadenyl tail (Supplementary Figure S1A).

Having validated *in vitro* translation (IVT) with the luciferase reporter system, we next attempted to optimize our platform for high throughput (HTP). Initial assays were scaled down from 50 to 20 μl and performed in 384-well plates. Measurements from five endpoints (at 0-4h post assembly of lysate with energy, amino acids and mRNA), each set up in triplicate, following the incubation at 37°C with shaking every 30 min in a plate reader were taken. For each time-point the luciferase reaction buffer was added directly into the test wells 10 minutes prior to measurement. An increase of detected luminescence demonstrated successful production of luciferase for at least 4h (Figure 1D).

We next aimed to investigate whether the continuous monitoring of the IVT might provide further information about the dynamics of parasite translation and drug dynamics over time. To this end, we explored generation of an mRNA reporter construct using conventional *P. pyralis* firefly luciferase versus an alternative green bioluminescence reporter from the click beetle *Pyrophorus plagiophthalamus* CBG99 ^15^. This was cloned into the pHLH construct replacing the *luc* expression cassette, generating a pH-CBG99-H expression vector with conserved flanking regulatory elements.

Customization of the luciferase reaction buffer (see MATERIALS AND METHODS), resulted in a 5fold increase of the signal to noise, which permitted a direct test between firefly and click beetle luciferase 5’hrp3 and 3’hrp2 flanked constructs to ascertain the optimal construct for signal intensity and continuous production of oxyluciferin, detectable with a plate reader. Of note, luminescence deriving from *CBG99* mRNA, but not *Luc*, produced a signal detectable with a peak at about 2.5 h post-set up using our customized luciferase reaction buffer, added at the start of the reaction/incubation time (Figure 2A). This suggested that the click beetle luciferase is either more stably produced over time or its activity more stable than that of firefly luciferase under our IVT assay conditions ^15^ (Figure 2A). We also explored the UTR-dependence of IVT using an alternative 5’ promoter element from the *P. falciparum erythrocyte binding antigen* (*eba*)-*175* (0.85 kb upstream of the ATG start site), a region previously validated for *in vitro* translation ^12^ and the human IVT compatible UTR pIRES, generating pEBA175-CBG99-H and pIRES-CBG99 respectively (Figure 2B).

**Figure 2.**
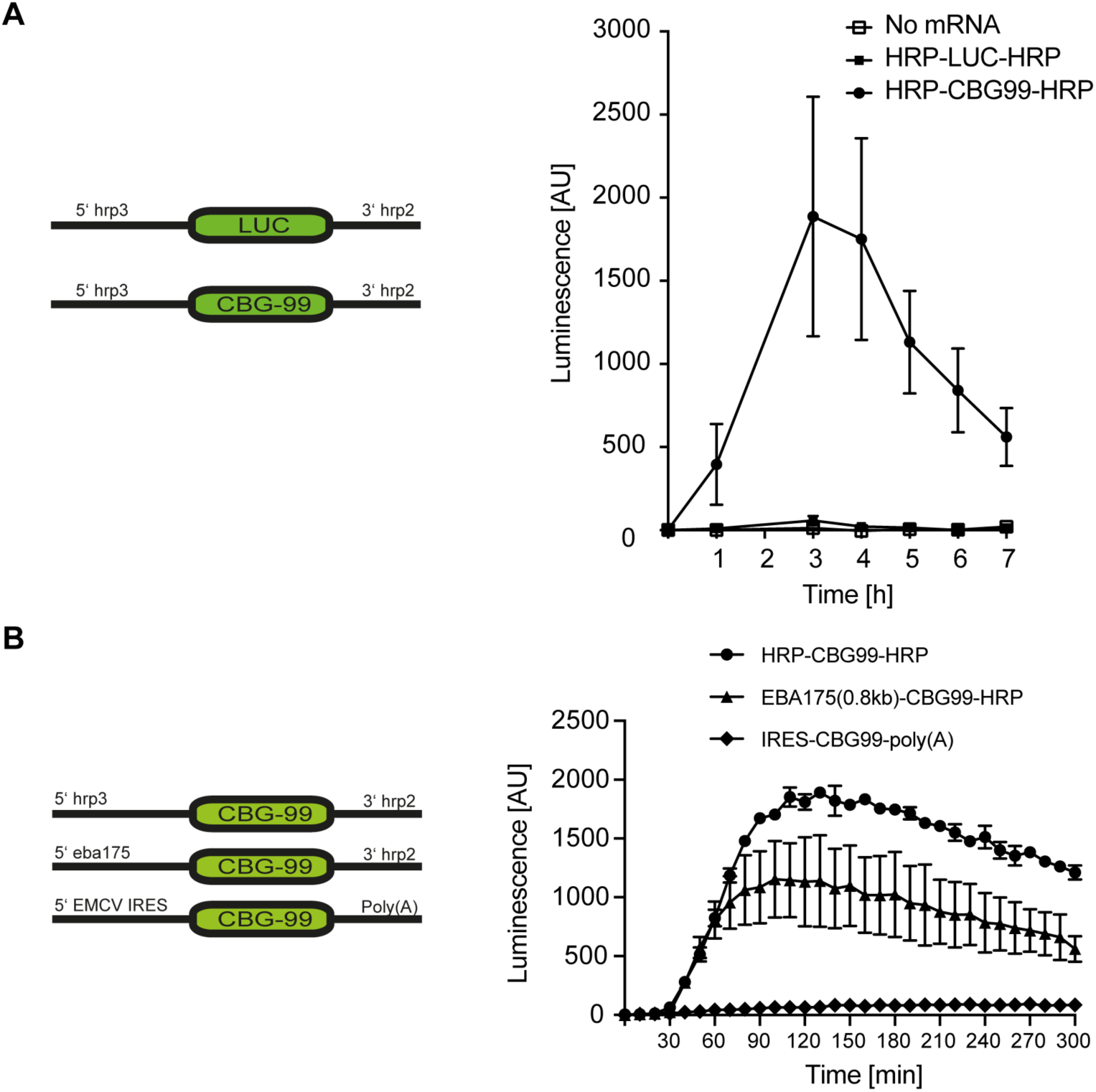
Optimization of the mRNA construct for continuous IVT. **A)** A time-course comparison of luminescence from *luc* and *cbg99* translated in lysate containing the luciferase reaction buffer from start of incubation to test stability of components in each assay. Left, schematics of the mRNA constructs used for *luc* and *cbg99* flanked by the histidine rich protein (hrp) 3 5’ UTR and hrp 2 3’ UTR. Right, the graph shows detected averages of triplicate assays with DMSO at indicated time points after start of the IVT assay/incubation (time point 0 h). Error bars indicate standard deviation. **B)** Comparison of *in vitro* translation levels of *cbg99* depending on the flanking UTRs. Left, schematics of the mRNA constructs used for *cbg99* flaked by either the histidine rich protein (hrp) 5’ and 3’ UTR, by a EBA-175 5’ UTR and hrp 3’UTR or the EMCV IRES 5’ UTR and a polyadenyl tail. Right, the graph shows detected averages of triplicate assays with DMSO at indicated time points after start of the IVT assay/incubation (time point 0 h). Error bars indicate standard deviation.

Comparison of the P*fhrp3* 5’ UTR with that from Pfeba-175, showed that whilst the *eba-175* 5’UTR is efficient in enabling IVT, as reported ^12^, especially when compared to an EMCV IRES using parasite lysate, the hrp3 5’UTR is considerably more efficient and reliable (smaller variation of signal between single assays) under our IVT conditions (Figure 2B). Minimal translation was seen using the pIRES-CBG99 vector. pH-CBG99-H mRNA was therefore selected for all further screening using our customized reaction buffer, with peak luminescence seen around 120 minutes.

To validate the ability of the plate-based luminescence assay for drug screening, we tested its sensitivity to known translation inhibitors including CHX and emetine (EME) (both of which target the parasite ribosome^16^) and halofuginone (HF) a direct target of the cytoplasmic *Plasmodium* protein translation enzyme prolyl-tRNA synthetase ^17^. Assayed in triplicate in the same experiment, each of these drugs showed inhibition of *in vitro* translation at nanomolar concentrations when measured at 2 hours (Figure 3A-C). Observed differences in the dynamics of the luminescence measured kinetically (time of onset, slope of increase, maximum and time-point at maximum measured luminescence) when assayed at 12 nM (Figure 3D) showed that each drug exhibited different residual activity. Whilst we have not explored this here, this may indicate that the assay could potentially determine not only translation inhibition during a screen (as an endpoint) but could be developed to interrogate potential mechanisms of action (as derived from the kinetics of activity) and stability of the drug when the whole profile of the curve of luciferase signal is considered. Work is ongoing towards validation of the concept. The dose dependency curves for the three drugs CHX, EME and HF (Figure 3A-C) were broadly consistent with experimentally determined cell culture IC50 values for emetine dihydrochloride hydrate (±47 nM) ^18^ and HF (±9 nM) ^17^ though less accurate for CHX, which is extremely potent in cellular assays (0.6 nM ^19^). This verified the sensitivity of the IVT assay towards a least two classes of translation inhibitors, demonstrating its utility in detecting compounds beyond those targeting only the ribosome.

**Figure 3.**
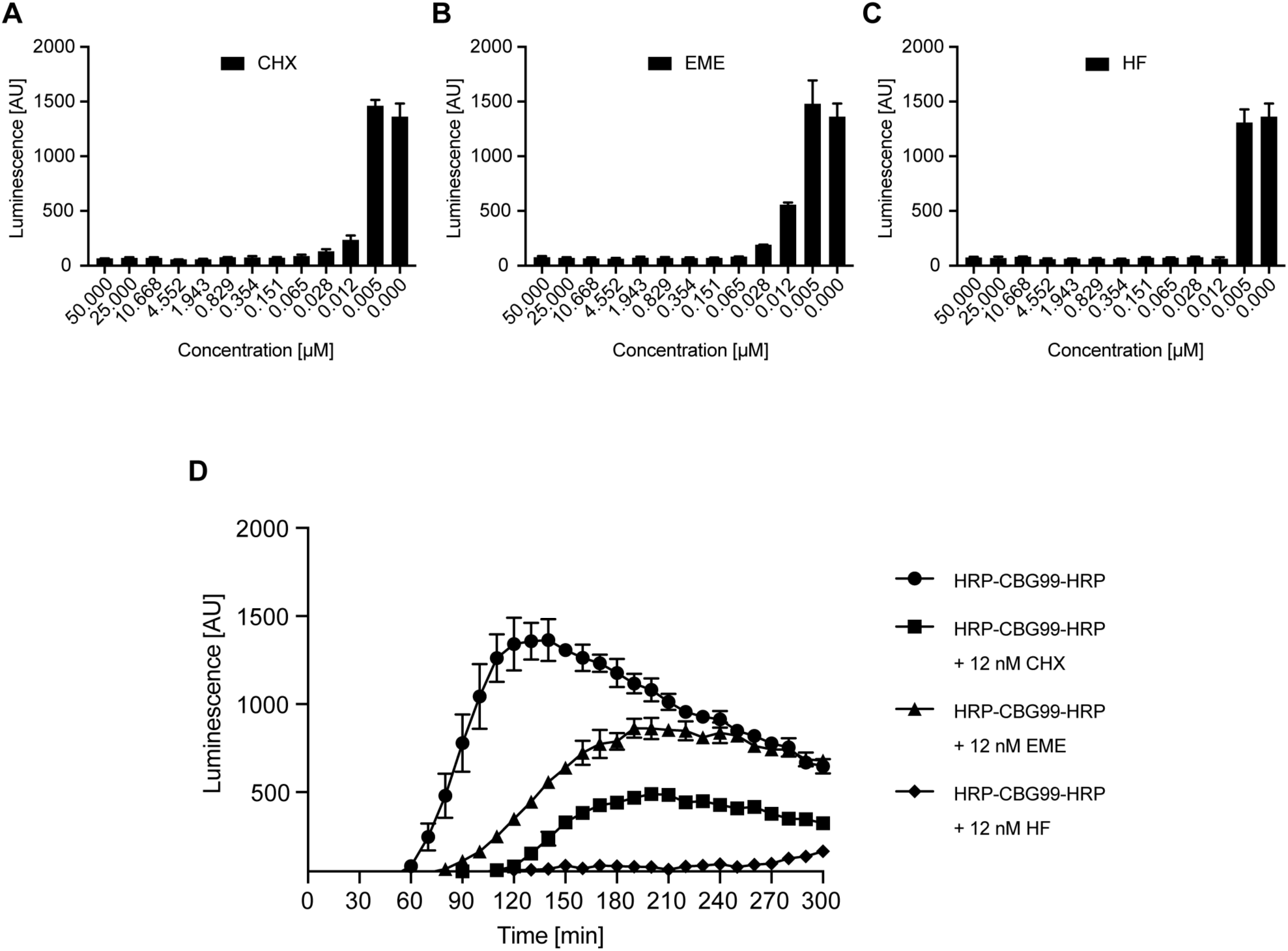
IVT Assay validation using known protein translation drugs. Bar charts show concentration-dependent inhibition of translation in presence of Cycloheximide, CHX (**A**), Emetine, EME (**B**) and Halofuginone, HF (**C**) respectively. Values arise from triplicate assays using a 12-point titration in the range of 50 μM to 0.005 μM measured at 120 minutes. **D**) Graph shows dynamic luminescence over time from triplicate assays containing either DMSO or drug at 12 nM. Error bars indicate standard deviation.

The Medicines for Malaria Venture (MMV) Malaria Box is a collection of 400 compounds representing families of structures identified as having activity against *P. falciparum* ^20^. Some of these were recently shown to affect parasite *in vitro* translation ^12^. Leading on from this we sought to validate the previously reported activity of candidate compounds compared to a third class of translation inhibitor, DDD107498, a current antimalarial in preclinical development that targets parasite protein translation *via* the translation elongation factor 2 (eEF2) ^8^. Contrary to published findings ^12^, measurement at 120 minutes did not show any substantial translation inhibition activity from the top 8 MMV compounds previously identified (Figure 4A, highlighted (blue labels) besides 8 further random MMV compounds (black)). Where activity was seen it was inconsistent between replicate runs (e.g. compound MMV006767, Figure 4A). It is unclear whether the poor activity seen in our screen is the result of use of a different parasite strain in our assay (3D7 versus W2mef), the buffer conditions used or differences in the 5’UTR. The validated translation inhibitor DDD107498, however, showed potent activity leading to a 73.5% inhibition of translation in our assay (Figure 4A). This further validates the breadth of activity detectable in our IVT assay, whilst suggesting that the Malaria Box is unlikely to have a potent translation inhibitor in its collection.

**Figure 4.**
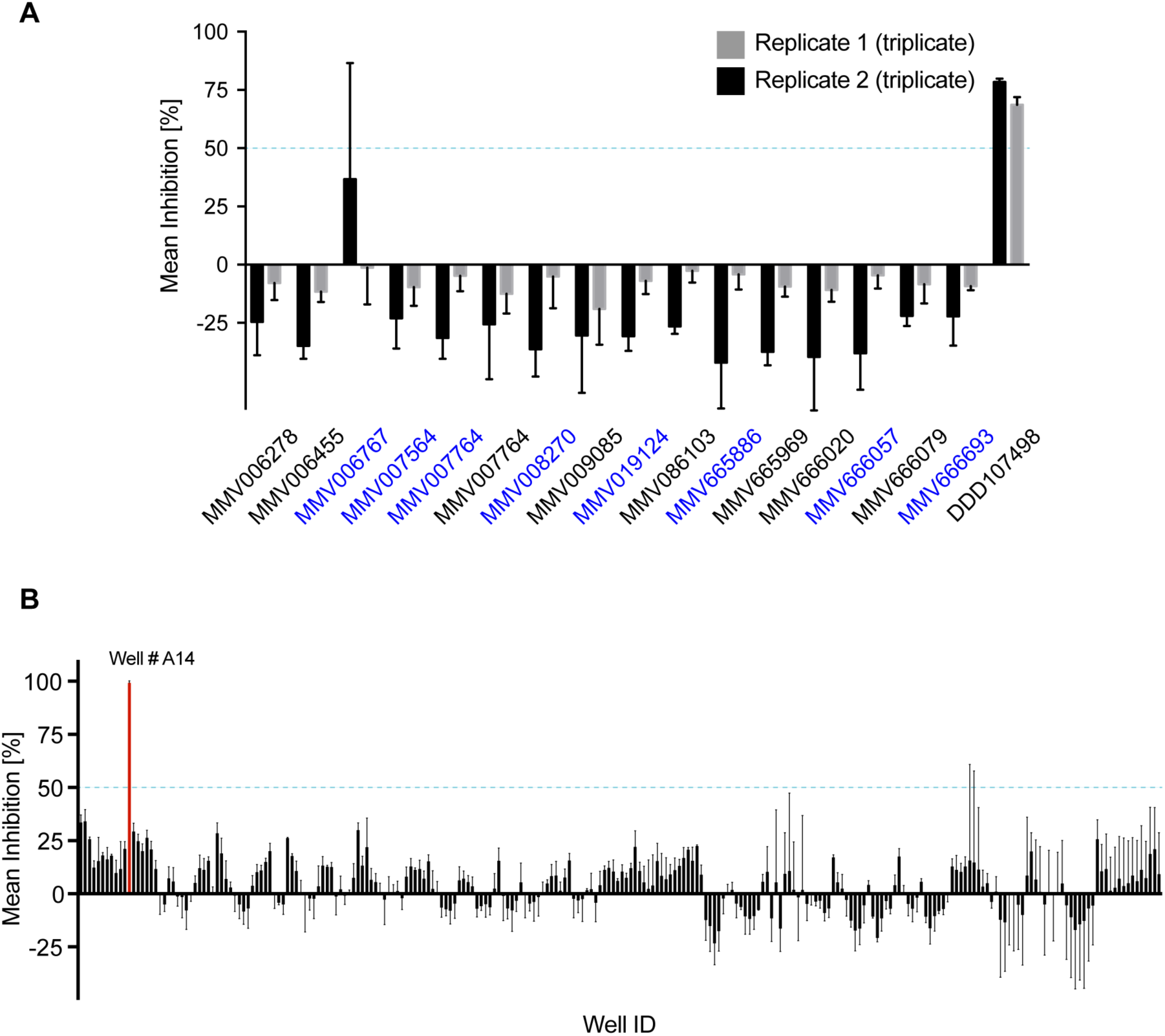
IVT Assay test of drug libraries. **A**) Screen of a selection of Medicines for Malaria Venture (MMV) Malaria box drugs compared with recently identified translation inhibitor DDD107498 ^8^. Bar graph shows the percentage of inhibition as mean of both of 2 independent assays (each in triplicate) with either 1 μM (final concentration) drug dissolved in DMSO measured at 120 minutes. Error bars indicate Standard Deviation. Drugs previously predicted to be active are labelled in blue. Blue horizontal lines indicate a threshold (50%) for drugs considered as hit. **B**) Blind screen of 247 selected compounds (enriched for potential antimicrobial activity) in HTP IVT assay in a 384-well plate format. Results shown in the bar graph are mean percentage inhibition from 3 independent assays (in singular form) from a 384-well plate containing 5 μM (final concentration) drug dissolved in DMSO measured at 120 minutes. Blue horizontal lines indicate a threshold (50%) for drugs considered as hits. For raw data values see Supplementary Data Table 1.

Finally, having verified the activity of our IVT platform for screening, we sought to undertake a blind HTP screen of untested compounds as validation for future screening of entire compound libraries. A proof-of-concept screen of 247 developmental antimicrobials in a 384-well plate format measured at 120 minutes was undertaken, with compounds selected from a ±2 million compound library (courtesy of GlaxoSmithKline) as having potential antimicrobial properties. Activity was generally very low with each compound tested, with few showing up to 30% inhibition of parasite protein translation (standardized for internal negative [DMSO] or positive [HF] controls) and none giving consistent inhibition across replicate plates (see Supplementary Data Table 1). One exception was a single compound identified as effectively reducing translation by ±99% (Figure 4B, compound A14). On un-blinding, this compound was revealed to be Puromycin, a drug known to inhibit protein translation, which has been previously shown to have nanomolar activity against *P. falciparum* ^21^.

**In conclusion**, we have optimized a cell free lysate for HTP IVT that reconstitutes the complete parasite translation machinery. The IVT assay is validated as effective in detecting compounds that inhibit ribosome activity directly (Emetine, Cycloheximide and Puromycin), ribosome co-factors (e.g. the recently identified compound DDD107498, targeting elongation factor elongation factor 2 (eEF2)) and tRNA synthetases (Halofuginone). We also describe the first blind screen for translation inhibitors using a small library of compounds enriched for anti-microbial activity, which although unable to find a new antimalarial hit successfully, found the only known translation inhibitor (Puromycin) in a blinded array of 247 compounds. This validates our HTP IVT platform, with potential to scale to several thousand compounds per week, for its ability to screen novel drug libraries for hits targeting malaria parasite protein translation. In continuous-read format, the IVT platform has the potential of providing immediate insights into the characteristics of each potential hit, where drugs of different mechanism may be revealed through the interrogation of their kinetic traces – a concept we are currently exploring. We believe the HTP IVT platform will provide a powerful addition to the antimalarial drug discovery pipeline.

## MATERIALS AND METHODS

### Vectors for *in vitro* mRNA production

Final plasmid sequences are available upon request. Maps are given in Supplementary Figure S1B.

#### pHLH-T7term

For efficient *in vitro* mRNA transcription, a T7 phage polymerase terminator region was introduced into the pHLH-1 downstream the 3’ hrp2-UTR by site directed mutagenesis (forward primer: CGCGCTTGGCGAATCATGGTCA, revers primer: GCTAGTTATTGCTCAGCGGCAATTAACCCTCACTAAAGGGAACAAAAG).

#### pH-CBG99-H

The firefly luciferase (*luc*) was replaced in the pHLH-T7term with the click beetle luciferase gene *cbg99*. Briefly, the *cbg99* expression cassette was PCR amplified from pCBG99 (Promega) using the forward primer <u>TTAATACAGTTATTTTAAAAAACC</u>ATGGTGAAGCGTGAGAAAAATGTCATC and reverse primer <u>TTTTTAATCTATTATTAAATAAGCTT</u>CTAACCGCCGGCCTTCTCCAA (homology regions are underlined) and pHLH-1 was restricted at its NsiI and HindIII sites prior to enzyme-dependent assembly cloning ^22^.

#### pEBA-CBG99-H

An 850 bp long region directly upstream of the EBA-175 start codon was PCR amplified from genomic DNA from 3D7 parasites using the forward primer <u>CGACGTTGTAAAACGACGGCCAGTGAATTCG</u>GAAGAAACAAGTGGTGTTCTAAAATATAATTA GC and the reverse primer <u>GATGACATTTTTCTCACGCTTCAC</u>CATTGTATGCACATTGAATATATTTATATGTTATTATC, the *cbg99* expression cassette was amplified from pCBG99 (Promega) using the forward primer ATGGTGAAGCGTGAGAAAAATGTCATC and reverse primer <u>TTTTTAATCTATTATTAAATAAGCTT</u>CTAACCGCCGGCCTTCTCCAA (homology regions are underlined) and pHLH-1 was restricted at its EcoRI and HindIII sites prior to enzyme-dependent assembly cloning.^22^

#### pIRES-Luc

The *luc* expression cassette from pHLH-1 was amplified by PCR using the forward primer, <u>CACGATGATAATATGGCCACC</u>ATGCATGAAGACGCCAAAAACATAAAG and the reverse primer <u>TCAGTGGTGGTGGTGGTGGTGCTC</u>CAATTTGGACTTTCCGCCCTTCTTG (homology regions are underlined) and introduced into a pT7CFE vector (Thermo Fisher Scientific) using the restriction sites NdeI and XhoI and enzyme-dependent assembly cloning, resulting in pT7CFE-luc.

#### pIRES-CBG99

The *cbg99* expression cassette from pCBG99 was amplified by PCR using the forward primer <u>GAAAAACACGATGATAATATGGCCA</u>CCATGGTGAAGCGTGAGAAAAATGT, and the reverse primer <u>CAGTGGTGGTGGTGGTGGTGCTCGAG</u>ACCGCCGGCCTTCTCCAACAAT (homology regions are underlined) and introduced into a pT7CFE vector (Thermo Fisher Scientific) using the restriction sites NdeI and XhoI and enzyme-dependent assembly cloning, resulting in pT7CFE-cbg99.

#### *Plasmodium falciparum* large scale culture

To prepare parasites for lysate production, 3-5 L of asexual *P. falciparum* parasites (3D7 strain) were grown in 225 cm^2^ flasks containing 300 mL complete RPMI (RPMI 1640 media supplemented with 25 mM HEPES (pH 7.4), 0.25% (w/v) sodium bicarbonate, 50 μg/ml hypoxanthine (Sigma-Aldrich), 25 ug/l gentamycin and 0.5% (w/v) Albumax II (Life Technologies)). Culture flasks were maintained at 2 % hematocrit blood (UK NHS Blood Transfusion Service), individually filled with ‘Malaria gas’ (3% (vol/vol) O_2_/5% CO_2_/92% N_2_ (BOC Special Gases)) and stored in a 37 °C incubator. Prior to seeding into large-scale flasks at 2% ring-stage parasitemia, parasites were synchronized with 5% sorbitol (Sigma Aldrich) to achieve high synchronicity ^23^. Media was exchanged daily until the culture reached 10-20% parasitemia. Cells were harvested at late trophozoite stage (approximately 30 h post-invasion) by centrifugation for 5 min at 800×*g* at room temperature (Eppendorf). Red blood cells were lysed in pellet wash buffer (PWB) containing 45 mM HEPES pH 7.4, 2 mM Mg(OAc)_2_, 100 mM KOAc, 250 mM Sucrose, 2 mM DTT, 20 U human placental RNase Inhibitor (2520), and 15 μM Leupeptin (L2884) supplemented with 0.075% (w/v) saponin (Sigma-Aldrich) for 5 min at room temperature. Lysed pellets were centrifuged at 4 °C 2800×*g* for 10 min and washed with ice-cold PWB. Wash steps were repeated until supernatant lost its red stain. Parasite pellets were either shock frozen and stored at −80°C for no longer than 1 month or immediately used for lysate production.

#### Production of *Plasmodium falciparum* IVT lysate

Washed pellets were re-suspended in 1-1.5 × pellet volumes of PWB supplemented with 1× EGTA-free protease inhibitor cocktail (Roche) and filled into a 45 mL pre-chilled nitrogen cavitation chamber (Parr Instrument Company) and incubated on ice at 1500 psi for 45 min. After release from the chamber the crude lysate was clarified by differential centrifugation, 15 min at 10,000×g followed by 15 min at 30,000x*g* at 4°C. As a measure for activity, the RNA content of the supernatant was checked using nanospectroscopy (Thermo Fisher Scientific). Lysates with a concentration above 1000 ng/μl were stored in aliquots of 100 μl at −80 °C.

#### Continuous *in vitro* translation and luciferase assay

For each IVT assay, several aliquots of different *P. falciparum* asexual stage lysate harvests were pooled to ensure consistency of activity. For dynamic assays validated with test drugs (Cycloheximide, Emetine and Halofuginone, Sigma-Aldrich) and screens, each reaction was assembled to a final volume of 20 μl. Each assay contained 25% lysate, 5% Amino acids (for final concentration 200 μM each, Sigma-Aldrich, stocks prepared according to Sun et al. ^24^), 5% energy and energy recovery system (final concentrations: 1.5 mM ATP, 0.15 mM GTP, 40 U/ml Creatine kinase (Sigma-Aldrich) and 4 mM Creatine phosphate (Fisher Scientific) in 40 mM HEPES), 5% additional components (final concentrations: 2% (w/w) PEG3000 (Sigma-Aldrich), 1 mM Spermidine (Sigma-Aldrich), 0.5 mM Folinic acid (Sigma-Aldrich), 200 μM Cystine and15 μM Leupeptin), 5% of supplemental HEPES, KOAC, Mg(OAC)_2_, DTT, RNase Inhibitor to equalize concentrations with PWB buffer, 4.5% 5’HRP3-CBG99-3’HRP2 mRNA (3500 ng/μl) *in vitro* transcribed from pH-CBG99-H (following ^14^), 0.5% DMSO or Drug dissolved in DMSO and, finally, 50% customized luciferase reaction buffer (45 mM HEPES (pH 7.4), 1 mM MgCl_2_, 1 mM ATP, 5 mM DTT, 1% (v/v) Triton-X, 10 mg/ml BSA, 1 × Reaction enhancer (Thermo Fisher Scientific), 1 mg/ml D-Luciferin). Provision of mRNA (following ^14^) involved incubation of 50-70 ng/μl of cDNA in a customized transcription buffer (40 mM HEPES, pH 7.4, 18 mM Mg(OAc)_2_, 5 mM each rNTPs (GE Healthcare Life Sciences), 2 mM Spermidine (Sigma), 40 mM DTT, 0.0025 U/μl Inorganic Phosphatase (Fermentas), 3 U/μl human placental RNase Inhibitor (Sigma), 10 U/μlT7-RNA Polymerase) for 100 min at 37 °C. 0.5 μl DNaseI (Fermentas) was added for the last 20 min. After dilution 1:1 (v/v) with RNase-free water, the mRNA was precipitated by addition of 0.5 × initial volume of 8 M LiCl followed by 30 min incubation on ice and subsequent centrifugation (20 min, Eppendorf 5424, full speed). The mRNA pellet was dissolved and precipitated again in presence of 0.3 M NH4OAc with 2.5 × volumes of absolute ethanol while kept at −80°C for 30min. The mixture was spun at full speed for 20 min (Eppendorf 5424), the mRNA pellet washed with 70% ethanol and dissolved in an appropriate volume of RNase-free water to a concentration of 3500 ng/μl shortly before experiments.

IVT reactions were carried out in low volume 384-well plates (Coring #3820 or Greiner BioOne #784101). DMSO and drugs were dispensed prior to assay assembly and plates were with sealed and stored at −20°C for up to 2 weeks. IVT components were dispensed using an ECHO 525 acoustic dispersion system (SynbiCITE, Imperial College London, UK) and incubated on drug for 15 min prior to addition of customized luciferase reaction buffer. Plates were then immediately transferred into a plate reader (Labtech, Tecan M200 Pro) and detection of luminescence in each well was carried out every 10 min over 5 h, while the sample was incubated at 37°C and shaken before every time-point.

For initial validation of *in vitro* translation of firefly luciferase and end-point measurements, assays were run in HEK cell and *Plasmodium falciparum* lysate 50 μl total assay volume containing 1% DMSO and either no mRNA or 5’EMCV IRES-Luc-3’Poly(A) mRNA *in vitro* transcribed from pT7CFE-Luc or 5’HRP3-Luc-3’HRP2 mRNA *in vitro* transcribed from pHLH-T7term respectively (Supplementary Figure S1A, Figure 1B-C). Assays were run over night (15 h) at 37°C with shaking every 30 min. Purchased luciferase reaction solution (Thermo Fisher Scientific) was added (1:1 v/v) using a multistep pipette (Eppendorf) 10 minutes before detection in the plate reader. For end-point-assays in 384-well plates, the reaction volume was scaled down to 20 μl lysate measurements were taken every hour over 4 h. Purchased luciferase reaction solution (Thermo Fisher Scientific) was added (1:1 v/v) using a multistep pipette (Eppendorf) 10 minutes before detection in the plate reader (Figure 1D).

To optimize continuous assay conditions, reactions were performed with either no mRNA or the following *in vitro* transcribed mRNAs: 5’HRP3-Luc-3’HRP2 from pHLH-T7term or 5’HRP3-CBG99-3’HRP2 from pH-CGB99-H. Customised luciferase reaction buffer (described above) was added using a multistep pipette immediately prior to the incubation at 37°C and start of measurement in the plate reader. Luminescence in each well was detected at time points 0 h, 1 h, 3 h, 4 h, 5 h, 6 h and 7 h (Figure 2A). To optimize 5’UTRs, reactions were performed with the following *in vitro* transcribed mRNAs: 5’HRP3-CBG99-3’HRP2 from pH-CBG99-H, 5’EBA175(0.8)-CBG99-3’HRP2 from pEBA-CBG99-H or 5’EMCV IRES-CBG99-3’Poly(A) mRNA *in vitro* transcribed from pT7CFE-CBG99 (Figure 2B).

#### Data Analysis

Results for each screen (Figure 4) were from either two, internal triplicated (MMV drugs) or three independent (antimicrobials) assays containing DMSO and HF as negative and positive controls respectively or drugs at varying concentrations. Inhibition (I [%]) was calculated from values at the time-point of maximum luminescence of the DMSO control (90-120 min after incubation start) normalized to the average control’s values at this time-point as: I = 100 – ((V_d_-μ_p_)/(μ_n_-μ_p_)*100), where V is the measured value, μ the is the mean, d the drug, p the positive controls and n the negative controls. Robustness (Z’ value) of plate based screens was calculated from the DMSO and HF controls using the formula Z’ = 1-((3 × (σ+σ_n_))/(|μ_p_-μ_n_|)), with σ as the standard deviation. Raw data of the tests are available upon request.

#### HEK cell large scale culture

0.5-1 L unmodified FreeStyle 293-F cells (Thermo Fisher Scientific) were grown in FreeStyle 293 Expression Medium in suspension to a density of 2 × 10^6^ cells/ml. Cells were pelleted for 15 min at 2.500 × g, 4 °C (Beckman Avanti centrifuge, JLA 10.500 Rotor) and washed by re-suspension in 5 × cell pellet volume PWB. After a subsequent centrifugation for 15 min at 2.500 × g, 4 °C (Eppendorf), cell pellets were either shock frozen and stored at −80°C for no longer than 6 months or immediately used for lysate production. Production of the IVT lysate, *in vitro* translation and luciferase assay were performed as described above with the exception that the mRNA for human IVT was transcribed from a pT7CFE vector (Thermo Fisher Scientific) so that the firefly luciferase was flanked by a EMCV IRES at the 5′ UTR and a poly adenyl tail.

#### Immunoblots

SDS-PAGE gels were run with 10 μl IVT lysate after 15 h incubation, incubated with 2 μl reducing sample buffer (95°C, 5 min), at 160 V for 45 min. Proteins were blotted onto a nitrocellulose membrane (Life Technologies) using an iBlot (Life Technologies) at program 0. The membrane was subsequently incubated in PBS containing 5% (w/v) skimmed milk powder for 1 h, before incubated with a mouse monoclonal anti-firefly luciferase antibody (Thermo Fisher Scientific) 1:2000 in PBS at 4°C overnight. After washing with PBS containing 1% (v/v) Tween-20, the membranes were incubated in horse radish peroxidase-conjugated anti-mouse (Sigma-Aldrich) antibody for 1 h at room temperature before washed and incubated in Amersham ECL Western Blotting reagent (GE Healthcare Life Sciences). Chemiluminescence was detected by exposing films (Carestream Kodak BioMax, Sigma-Aldrich) for 10 s and developing in an automated developer.

## ACKNOWLEDGMENTS

We thank Vida Ahyong and Joseph Derisi (UCSF) for generously sharing pre-publication data and methodological insights into *Plasmodium in vitro* translation. We thank Fernando Sanchez-Roman Teran (Imperial College London) for help with optimization of *Plasmodium falciparum* large-scale culture, Maria G. Gomez-Lorenzo, Francisco Javier Gamo and Luis Ballell-Pages (GlaxoSmithKline) and Michael Delves (Imperial College London) for providing test drugs, experimental advice and productive discussions regarding HTP screening and data analysis. We thank David McClymont (SynbiCITE) for thoughtful discussions about assay design and providing access to the ECHO252 system for lysate printing. We also thank Niklas Rehmert and Marko Storch (Imperial College London) for assistance in the early stages of this project and the UK NHS Blood Transfusion Service for generous provision of red blood cells. The research was directly supported by the Imperial Confidence in Concept Scheme, NIHR Imperial Biomedical Research Centre / Imperial Innovations Therapeutic Primer Fund & NIHR BRC at the Royal Marsden and the Institute of Cancer Research Funds 2014 (P51477) (J.B. & K.R.W.) and a Wellcome Pathfinder Award (105686) (J.B.). J.B. is supported by a Wellcome Investigator Award (100993/Z/13/Z).

## AUTHORSHIP CONTRIBUTIONS

H.B., H.M., K.R.W., and J.B. designed all experiments; H.B., H.M., M.L, and J.J.H. performed experiments; All authors contributed to manuscript preparation; H.B. and J.B. wrote the manuscript.

## REFERENCES

1 Bhatt, S. et al. The effect of malaria control on Plasmodium falciparum in Africa between 2000 and 2015. Nature 526, 207-211, doi:10.1038/nature15535 (2015).

2 WHO. World Malaria Report. (WHO, 2016).

3 Partnership, R. C. T. Efficacy and safety of RTS,S/AS01 malaria vaccine with or without a booster dose in infants and children in Africa: final results of a phase 3, individually randomised, controlled trial. Lancet 386, 31-45, doi:10.1016/S0140-6736(15)60721-8 (2015).

4 Ashley, E. A. et al. Spread of artemisinin resistance in Plasmodium falciparum malaria. N Engl J Med 371, 411-423, doi:10.1056/NEJMoa1314981 (2014).

5 Menard, D. et al. A Worldwide Map of Plasmodium falciparum K13-Propeller Polymorphisms. N Engl J Med 374, 2453-2464, doi:10.1056/NEJMoa1513137 (2016).

6 Wells, T. N., Hooft van Huijsduijnen, R. & Van Voorhis, W. C. Malaria medicines: a glass half full? Nat Rev Drug Discov 14, 424-442, doi:10.1038/nrd4573 (2015).

7 Gamo, F.-J. et al. Thousands of chemical starting points for antimalarial lead identification. Nature 465, 305-310, doi:10.1038/nature09107 (2010).

8 Baragana, B. et al. A novel multiple-stage antimalarial agent that inhibits protein synthesis. Nature 522, 315-320, doi:10.1038/nature14451 (2015).

9 Kato, N. et al. Diversity-oriented synthesis yields novel multistage antimalarial inhibitors. Nature 538, 344-349, doi:10.1038/nature19804 (2016).

10 Wilson, D. N. Ribosome-targeting antibiotics and mechanisms of bacterial resistance. Nat Rev Microbiol 12, 35-48, doi:10.1038/nrmicro3155 (2014).

11 Ferreras, A., Triana, L., Correia, H., Sanchez, E. & Herrera, F. An in vitro system from Plasmodium falciparum active in endogenous mRNA translation. Mem Inst Oswaldo Cruz 95, 231-235 (2000).

12 Ahyong, V. et al. Identification of Plasmodium falciparum specific translation inhibitors from the MMV Malaria Box using a high throughput in vitro translation screen. Malar J 15, 173, doi: 10.1186/s12936-016-1231-8 (2016).

13 Wu, Y., Sifri, C. D., Lei, H. H., Su, X. Z. & Wellems, T. E. Transfection of Plasmodium falciparum within human red blood cells. Proc Natl Acad Sci U S A 92, 973-977 (1995).

14 Mureev, S., Kovtun, O., Nguyen, U. T. & Alexandrov, K. Species-independent translational leaders facilitate cell-free expression. Nat Biotechnol 27, 747-752, doi:10.1038/nbt.1556 (2009).

15 Wood, K. V., Lam, Y. A. & McElroy, W. D. Introduction to beetle luciferases and their applications. J Biolumin Chemilumin 4, 289-301, doi:10.1002/bio.1170040141 (1989).

16 Wong, W. et al. Cryo-EM structure of the Plasmodium falciparum 80S ribosome bound to the anti-protozoan drug emetine. eLife, e03080, doi:10.7554/eLife.03080 (2014).

17 Jain, V. et al. Structure of Prolyl-tRNA Synthetase-Halofuginone Complex Provides Basis for Development of Drugs against Malaria and Toxoplasmosis. Structure 23, 819-829, doi:10.1016/j.str.2015.02.011 (2015).

18 Matthews, H., Usman-Idris, M., Khan, F., Read, M. & Nirmalan, N. Drug repositioning as a route to anti-malarial drug discovery: preliminary investigation of the in vitro anti-malarial efficacy of emetine dihydrochloride hydrate. Malar J 12, 359, doi:10.1186/1475-2875-12-359 (2013).

19 Gershon, P. D. & Howells, R. E. Mitochondrial protein synthesis in Plasmodium falciparum. Mol Biochem Parasitol 18, 37-43 (1986).

20 Van Voorhis, W. C. et al. Open Source Drug Discovery with the Malaria Box Compound Collection for Neglected Diseases and Beyond. PLoS Pathog 12, e1005763, doi:10.1371/journal.ppat.1005763 (2016).

21 Mamoun, C. B., Gluzman, I. Y., Goyard, S., Beverley, S. M. & Goldberg, D. E. A set of independent selectable markers for transfection of the human malaria parasite Plasmodium falciparum. Proc Natl Acad Sci U S A 96, 8716-8720 (1999).

22 Gibson, D. G. et al. Enzymatic assembly of DNA molecules up to several hundred kilobases. Nat Methods 6, 343-345, doi:10.1038/nmeth.1318 (2009).

23 Lambros, C. & Vanderberg, J. P. Synchronization of Plasmodium falciparum erythrocytic stages in culture. J Parasitol 65, 418-420 (1979).

24 Sun, Z. Z. et al. Protocols for implementing an Escherichia coli based TX-TL cell-free expression system for synthetic biology. J Vis Exp, e50762, doi:10.3791/50762 (2013).

